## Supplementary Table 1 for "R2DT: computational framework for template-based RNA secondary structure visualisation across non-coding RNA types"

### Supplementary Information

**Supplementary Table 1**. Newly developed rRNA templates using experimental data (PDB and chain identifiers)

| **rRNA type** | **Species** | **Clade** | **Supporting experimental data** |
| --- | --- | --- | --- |
| LSU | *Mycobacterium tuberculosis* | Bacteria (Actinobacteria) | 5V7Q:A |
| LSU | *Staphylococcus aureus* | Bacteria (Firmicutes) | 4WF9:X |
| LSU | *Tetrahymena thermophila* | Eukaryota | 4V8P:D1+C2 |
| LSU | *Bacillus subtilis* | Bacteria (Firmicutes) | 5NJT:U |
| LSU | *Homo sapiens* mitochondria | Eukaryota | 5OOL:A |
| LSU | *Homo sapiens* | Eukaryota | 6EK0:L5+L8 |
| LSU | *Dictyostelium discoideum* | Eukaryota | 5AN9:N |
| LSU | *Thermus thermophilus* | Bacteria (Deinococcus-Thermus) | 4Y4O:2A |
| LSU | *Haloarcula marismortui* | Archaea | 4V9F:0 |
| LSU | *Mycobacterium smegmatis* | Bacteria (Actinobacteria) | 5O60:A |
| LSU | *Plasmodium falciparum* | Eukaryota | 3J79:A+C |
| LSU | *Saccharomyces cerevisiae* | Eukaryota | 5TBW:1+4 |
| LSU | *Spinacia oleracea* chloroplast | Eukaryota | 6ERI:AA |
| LSU | *Escherichia coli* | Bacteria (Gammaproteobacteria) | 5J7L:DA |
| LSU | *Deinococcus radiodurans* | Bacteria (Deinococcus-Thermus) | 4IOA:X |
| LSU | *Drosophila melanogaster* | Eukaryota | 4V6W:A5 |
| LSU | *Kluyveromyces lactis* | Eukaryota | 4V91:1+4 |
| LSU | Lokiarcheaota F3H4 | Archaea | Based on ^1^ |
| LSU | Lokiarcheaota GC1475 | Archaea | Based on ^1^ |
| LSU | *Tetrahymena thermophila* mitochondria | Eukaryota | Based on ^1^ |
| SSU | *Homo sapiens* | Eukaryota | 4V6X:B2 |
| SSU | *Thermus thermophilus* | Bacteria (Deinococcus-Thermus) | 4V51:AA |
| SSU | *Pyrococcus furiosus* | Archaea | 4V6U:A2 |
| SSU | *Drosophila melanogaster* | Eukaryota | 4V6W:B2 |
| SSU | *Saccharomyces cerevisiae* | Eukaryota | 4V88:A2 |
| SSU | *Trypanosoma brucei* | Eukaryota | 4V8M:AA |
| SSU | *Escherichia coli* | Bacteria (Gammaproteobacteria) | 4V9D:AA |

Supplementary Table 2. Summary of template to diagram similarity by phylogenetic rank for RefSeq rRNA sequences.

| **Taxonomic Rank** | **Number of nucleotides positioned exactly as in template** | **Number of nucleotides inserted compared to template** | **Number of nucleotides requiring repositioning compared to template** | **Total number of displayed nucleotides** | **Number of diagrams** |
| --- | --- | --- | --- | --- | --- |
| species | 259,034 (97.%) | 4,175 (1.6%) | 3,896 (1.5%) | 267,105 | 147 (.6%) |
| genus | 538,357 (98.%) | 5,759 (1.%) | 5,332 (1.%) | 549,448 | 384 (1.6%) |
| family | 868,585 (98.%) | 9,279 (1.%) | 8,629 (1.%) | 886,493 | 558 (2.3%) |
| order | 1,390,705 (98.%) | 15,281 (1.1%) | 13,761 (1.%) | 1,419,747 | 860 (3.6%) |
| class | 6,264,956 (97.5%) | 68,719 (1.1%) | 89,065 (1.4%) | 6,422,740 | 3,844 (16.1%) |
| phylum | 7,094,429 (97.3%) | 104,029 (1.4%) | 91,687 (1.3%) | 7,290,145 | 4,780 (20.%) |
| kingdom | 20,172,007 (95.4%) | 557,422 (2.6%) | 420,121 (2.%) | 21,149,550 | 13,224 (55.5%) |
| superkingdom | 28,360 (94.2%) | 757 (2.5%) | 997 (3.3%) | 30,114 | 8 (.%) |
| root | 20,410 (37.5%) | 838 (1.5%) | 33,142 (60.9%) | 54,390 | 38 (.2%) |
| **Total** | **36,636,843 (96.2%)** | **766,259 (2.%)** | **666,630 (1.8%)** | **38,069,732** | **23,843** |

### References

1.	Penev, P. I. *et al.* Supersized ribosomal RNA expansion segments in Asgard archaea. doi:[10.1101/2019.12.25.888164](http://dx.doi.org/10.1101/2019.12.25.888164).
